# Sensory domain of the cell cycle kinase CckA in *Caulobacter crescentus* regulates the differential DNA binding activity of the master regulator CtrA

**DOI:** 10.1101/300178

**Authors:** Sharath Narayanan, Lokesh Kumar, Sunish Kumar Radhakrishnan

## Abstract

Sophisticated signaling mechanisms allow bacterial cells to cope with environmental and intracellular challenges. Activation of specific pathways facilitates the cells to overcome cellular damage and thereby warrant integrity. Here we demonstrate the pliability of the CckA-CtrA two component signaling system in the freshwater bacterium, *Caulobacter crescentus*. Our forward genetic screen to analyse suppressor mutations that can negate the chromosome segregation block induced by the topoisomerase IV inhibitor, NstA, yielded various point mutations in the cell cycle histidine kinase, CckA. Notably, we identified a point mutation in the PAS-B domain of CckA, which resulted in increased levels of phosphorylated CtrA (CtrA∼P), the master cell cycle regulator. Surprisingly, this increase in CtrA∼P levels did not translate into a genome-wide increase in the DNA occupancy of CtrA, but specifically enriched its affinity to the chromosomal origin of replication, C_*ori*_, and a very small sub-set of CtrA regulated promoters. We show that through this enhanced binding of CtrA to the C_*ori*_, cells are able to overcome the toxic defects rendered by stable NstA through a possible slow down in the chromosome cycle. Taken together, our work opens up an unexplored and intriguing aspect of the CckA-CtrA signal transduction pathway. The distinctive DNA binding nature of CtrA and its regulation by CckA might also be crucial for pathogenesis because of the highly conserved nature of CckA-CtrA pathway in alphaproteobacteria.

## Introduction

Bacteria harbor robust signaling mechanisms, to respond to numerous environmental challenges both inside and outside the cell. Exquisitely fine tuned regulatory cascades in bacteria impart their effect at a precise spatio-temporal scale to bring about specific morphological and functional programs, in response to the changes in the internal or external milieu. The aquatic α-proteobacterium, *Caulobacter crescentus* (henceforth *Caulobacter*), has emerged as a powerful model organism for studying the complex signaling mechanisms that control cell cycle and development in response to environmental cues. During its cell cycle, *Caulobacter* undergoes asymmetric division to produce progenies with distinct developmental fates. One of the daughter cells, the swarmer cell, acquires a dispersal fate wherein its motility is assisted by the polar flagellum and a tuft of pili (1,2). In contrast, the stalked daughter cell acquires a sedentary fate and is in an S-phase-like state capable of replicating its chromosome and proliferating by cytokinesis (Figure 1A) (3,4). The G1-like swarmer cell has to terminally differentiate into a stalked cell to enter into the proliferative phase. This G1 to S-like transition is marked by the shedding of the flagellum, retraction of the pili, and production of a stalk at the same cell pole (Figure 1A).

**Figure 1.**
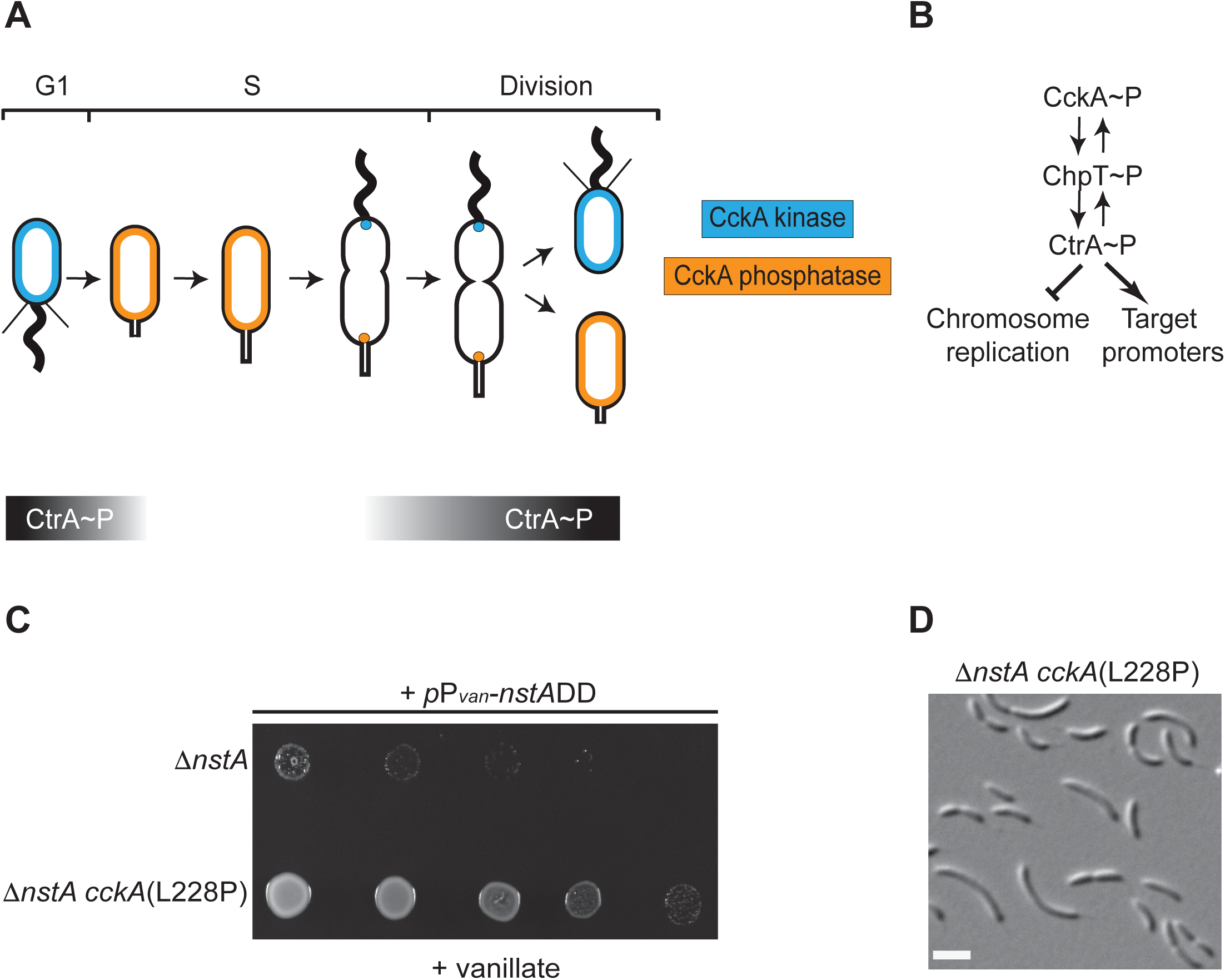
Cell cycle regulation in *Caulobacter crescentus* by the CckA-CtrA pathway. (A) Schematic representation of the dual switching of CckA between kinase mode (blue) and phosphatase mode (orange) in the swarmer and stalked cell compartments, respectively. The graded bars indicate the time during which CtrA (black) is present during the cell cycle. (B) The bidirectional flow of phosphate between CckA, ChpT and CtrA. In the swarmer cells, CckA transfers the phosphate group to the phosphotranferase, ChpT, which further donate phosphate group to CtrA. Thus, the phosphorylated form of CtrA (CtrA∼P) represent the active form, wherein CtrA can bind to various target promoters of several cell regulated genes, as well as repress the initiation of chromosome replication. (C) Growth of ∆*nstA*, and the suppressor mutant, ∆*nstA cckA*(L228P), upon *nstA*DD overexpression. Cells, as indicated, were diluted five folds and spotted on media containing 0.5 mM vanillate. (D) Differential interference contrast (DIC) image of the suppressor mutant, ∆*nstA cckA*(L228P). Scale bar: 2µm.

In the swarmer cells, the master transcriptional regulator, CtrA, inhibits the DNA replication. The *Caulobacter* origin of replication, C_*ori*_, is bound by CtrA, which prevents replisome formation in the swarmer cells (5). Concurrent with the swarmer to stalked cell transition, CtrA is degraded by proteolysis and thus facilitating the binding of DnaA, the replication initiator, to the C_*ori*_ triggering chromosome replication.(6). Apart from blocking DNA replication initiation, CtrA also serve as a transcription factor to drive the expression of numerous developmentally important genes in a cell cycle dependent manner (7).

The differential activity of CtrA in swarmer and stalked cells is of paramount significance for generating different cell fates. Multiple levels of regulation involving control at the level of synthesis, stability, and activity exist for the regulation of CtrA during cell cycle (8,9). The phosphorylated form of CtrA (CtrA∼P) represents the active form that binds to DNA (10). The phosphorylation of CtrA is catalyzed by an essential hybrid cell cycle histidine kinase/phosphatase, CckA, which phosphorylates CtrA through the single domain histidine phosphotransferase, ChpT (11–14). In the swarmer, and pre-divisional cells, the kinase activity of CckA ensures the abundance of active CtrA∼P, while in the stalked cell compartment, the phosphatase activity of CckA is predominant ensuring the dephosphorylation, and degradation, of CtrA (Figures 1A and 1B) (15).The bifunctional nature of CckA is governed by the second messenger, cyclic-di-guanylate (c-di-GMP). Binding of c-di-GMP to CckA causes an inhibition of its kinase activity and trigger the phosphatase activity. The swarmer to stalked cell transition is accompanied by an increase in the levels of c-di-GMP, which causes the transition of CckA from the kinase to the phosphatase mode (16,17).

Recent evidences have shown that in addition to developmental regulatory proteins such as CtrA, the cell cycle progression in *Caulobacter* is controlled by a cytoplasmic redox fluctuation (18). We have shown that a redox dependent regulator, NstA, whose activation is coupled to the cytoplasmic redox state, inhibits the DNA decatenation activity of topoisomerase IV (Topo IV) during the early stages of cell cycle (18). Apart from the cytoplasmic redox control of NstA activity, additional layers of regulation for NstA exist at the level of transcription by the transcription factors, GcrA and CcrM, and at the level of protein abundance by the ClpXP protease. A stable version of NstA, NstADD, is resistant to protein degradation by ClpXP. Overproduction of NstADD from an inducible promoter induces lethality in *Caulobacter* (18). In this study, we wished to investigate the regulatory networks that possibly fine-tune NstA activity *in vivo*. Towards this, we exploited the lethality induced by NstADD, to conduct an unbiased forward genetic screen to analyze extragenic suppressor mutation(s) that can negate NstADD toxicity. Strikingly, through this screen we have identified suppressor mutations in *cckA* that influences the DNA binding activity of CtrA in a distinctive manner. We show that the CckA(L228P) mutation though enhances the CtrA∼P levels, does not universally increase the binding of CtrA on a genome-wide scale. Rather, we found that the CckA(L228P) mutation specifically increases the binding of CtrA∼P at the C_*ori*_ and a very small sub-set of CtrA dependent promoters. Finally, we show that the enhanced binding of CtrA to the C_*ori*_ rescues the toxicity caused by NstADD by possibly slowing down the chromosome replication process to compensate for the slowed down segregation caused by the inhibitory effects of NstADD on the Topo IV.

## Results

### Suppressor mutations in CckA alleviate NstADD toxicity

Overproduction of the cell-cycle stable form of the Topo IV inhibitor, NstADD, leads to fitness defect and impaired chromosome segregation in *Caulobacter* (18). To unearth the signaling network that regulate NstA, we exploited the lethality induced by NstADD overproduction in a genetic screen to identify extragenic suppressors that could tolerate NstADD toxicity (See Materials and Methods). Strikingly, whole-genome sequencing revealed that all nine extragenic suppressors harbored a mutation in the gene encoding the cell cycle histidine kinase, *cckA* (Supplementary Figure S1C). The suppressor mutations were CckA-(L228P), (A317V), (D364G), (R356C), (F392L), (F493C) and (F496C) (Figure 1C, Supplementary Figures S1B, S2A and S2B). Interestingly, all the point mutations in *cckA*, except the L228P substitution, were located either in the histidine kinase domain or the ATP binding domain of the CckA protein (Supplementary Figure S1C). Remarkably the L228P mutation, which mapped to the PAS-B domain in CckA, rendered developmental defects such as cell filamentation in the *WT* and the *∆nstA* mutant (Figure 1D, Supplementary Figure S2A). To confirm that it is the L228P mutation in *cckA* that conferred resistance to NstADD toxicity, we backcrossed the *cckA*(L228P) mutation into *WT* and *∆nstA*. The backcrossed cells were indeed able to tolerate the NstADD overexpression (Supplementary Figure S1A). The fact that the CckA(L228P) mutation was not in the kinase or the ATP binding domain of CckA, prompted us to investigate further the mechanism by which the *cckA*(L228P) mutant induced the developmental defects and negated the chromosome segregation defect attributed by NstADD in *Caulobacter*.

### CckA(L228P) mutation leads to increased CtrA∼P levels but not increase in CtrA binding or activity

The cell cycle histidine kinase, CckA, is the primary kinase that activates the master cell cycle transcriptional regulator, CtrA, by phosphorylation (10). The phosphorylated form of CtrA (CtrA∼P) binds efficiently to its target promoters on the chromosome regulating transcription (10,11,19). Therefore, we decided to investigate if the L228P mutation in the PAS-B domain of CckA could affect CtrA∼P levels. Interestingly, *in vivo* phosphorylation analysis revealed that the relative levels of CtrA∼P, compared to total CtrA, was two fold higher in the *cckA*(L228P) mutant than the wild type cells (Figure 2A).

**Figure 2.**
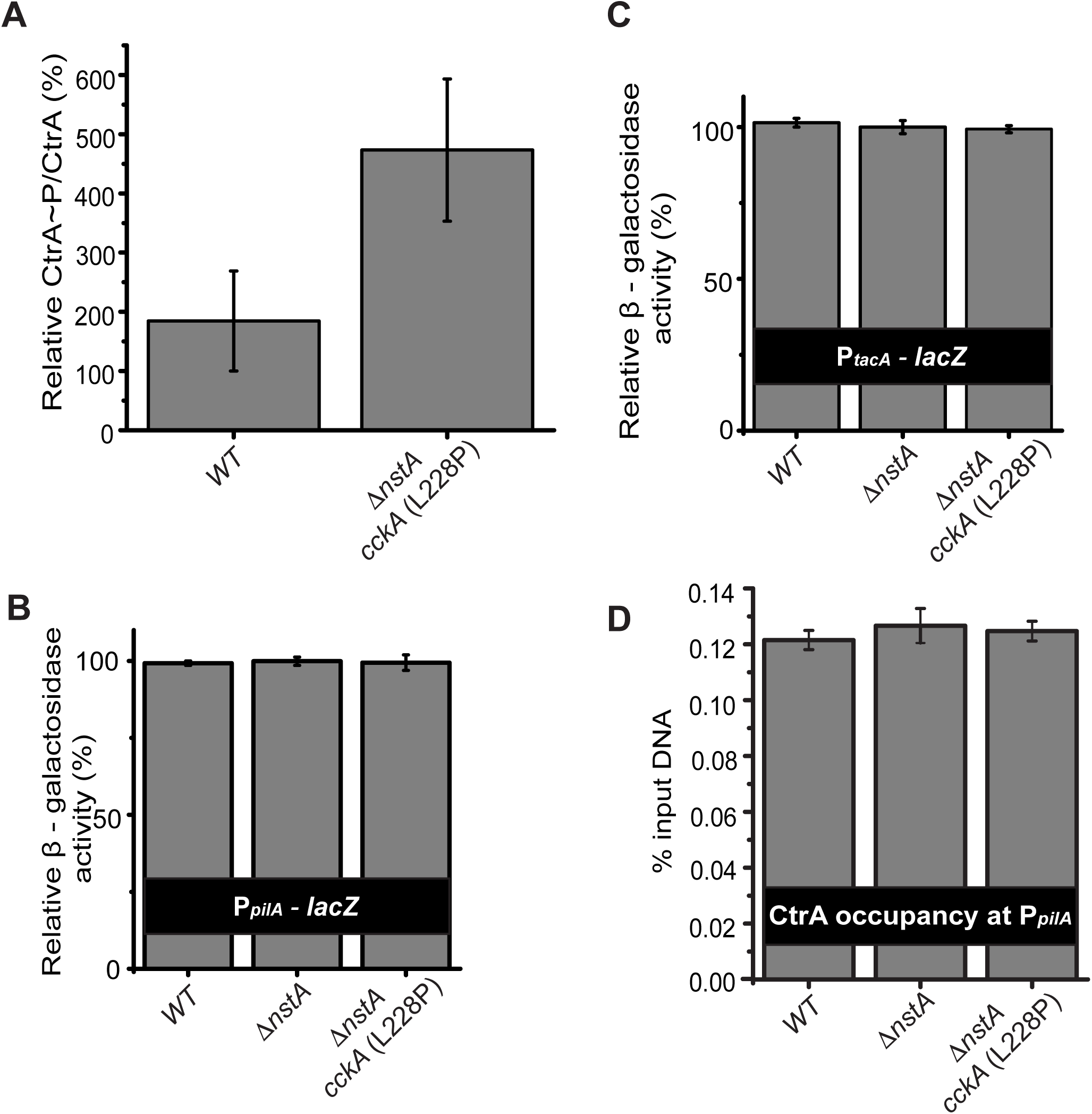
Effect of CckA(L228P) mutation on CtrA. (A) *In vivo* phosphorylation experiment denoting CtrA∼P/CtrA levels in *WT* and ∆*nstA cckA*(L228P) mutants. The relative β-galactosidase activity (in percentage) of (B) the P_*tacA*_-*lacZ* and (C) the P_*pilA-*_*lacZ* reporters in *WT*, ∆*nstA* and ∆*nstA cckA*(L228P) cells. (D) Data of qChIP analysis showing the CtrA occupancy at the promoter region of *pilA* (P_pilA_) in *WT*, ∆*nstA* and ∆*nstA cckA*(L228P) cells. The values ± SE, represented in A, B, C and D are the average of at least three independent experiments.

It has been shown that increase in CtrA∼P levels could result in elevated levels of CtrA binding to its target promoters (20,21). Therefore, we decided to analyze the binding of CtrA∼P on well-established CtrA-dependent promoters such as *pilA*, *tacA* and *kidO* (22–24), by quantitative chromatin immunoprecipitation (qChIP) analysis using CtrA specific antibodies. Surprisingly, our qChIP experiments revealed that the binding of CtrA to the P_*pilA*_, P_*kidO*_ and P_*tacA*_ promoters were not significantly different in the *cckA*(L228P) mutant when compared to wild-type (Figure 2D, Supplementary Figure S3A, B). Further, β-galactosidase (LacZ)-based promoter-probe assays using the promoters of *pilA* and *tacA* in *WT*, ∆*nstA* and ∆*nstA cckA*(L228P) mutant backgrounds, revealed that there was no measurable differences in the P_*pilA*_ and P_*tacA*_ promoter activities in the *cckA*(L228P) mutant (Figures 2B and 2C).

Collectively, these results indicated that the *cckA*(L228P) mutation increased CtrA∼P levels. Nevertheless, this surge in CtrA∼P levels did not result in increased binding of CtrA, or elevated promoter activity, on at least the G1-specific promoters of CtrA such as P_*kidO*_, P_*pilA*_ and P_*tacA*_. Furthermore, these observations opened up the possibility that CckA, apart from its kinase activity, may be influencing the DNA-binding activity of CtrA, only at specific promoter regions in the *Caulobacter* genome.

### CckA influences the promoter specific binding of CtrA

Next we decided to investigate if the absence of difference in binding of CtrA, despite increased CtrA∼P levels in the *cckA*(L228P) mutant, is specific to a subset of CtrA dependent promoters. Towards this, we performed chromatin immunoprecipitation followed by deep sequencing (ChIP-seq) to analyze the CtrA occupancy on its target promoters in the *cckA* and *cckA*(L228P) mutant backgrounds on a genome-wide scale. From the ChIP-seq analyses it was evident that the *cckA*(L228P) mutation enhanced the CtrA occupancy at the promoter regions of target genes whose transcripts peaked at late S-phase, including the Class II flagellar genes such as *pleA, fliQ, fliL, fliM*, *fliJ* and *fliI* (25), pilus secretion genes, *cpaA* and *cpaB* (26), flagellar regulatory genes, *flbT* and *flbA* (27), and the chemotaxis genes, *motA* and *motB* (28) (Figures 3 A, C, D and Supplementary Dataset 1). To corroborate if this increase in binding of CtrA to these promoters resulted in an increased promoter activity, we analyzed the activity of the *fliM* promoter (P_*fliM*_) and the *flbT* promoter (P_*flbT*_) using LacZ reporter fusions to these promoters (P_*fliM*_-*lacZ* and P_*flbT*_-*lacZ*). Our analyses showed that indeed the activities of P_*fliM*_ and P_*flbT*_ were increased in the *cckA*(L228P) mutant background commensurate with the increase in binding of CtrA to these promoters (Supplementary Figure S4A, B). Further, the qChIP experiments confirmed the enhanced binding of CtrA at the promoter region of *flbT*, in the ∆*nstA cckA* (L228P) background when compared to the WT or ∆*nstA* (Supplementary Figure S3C).

**Figure 3.**
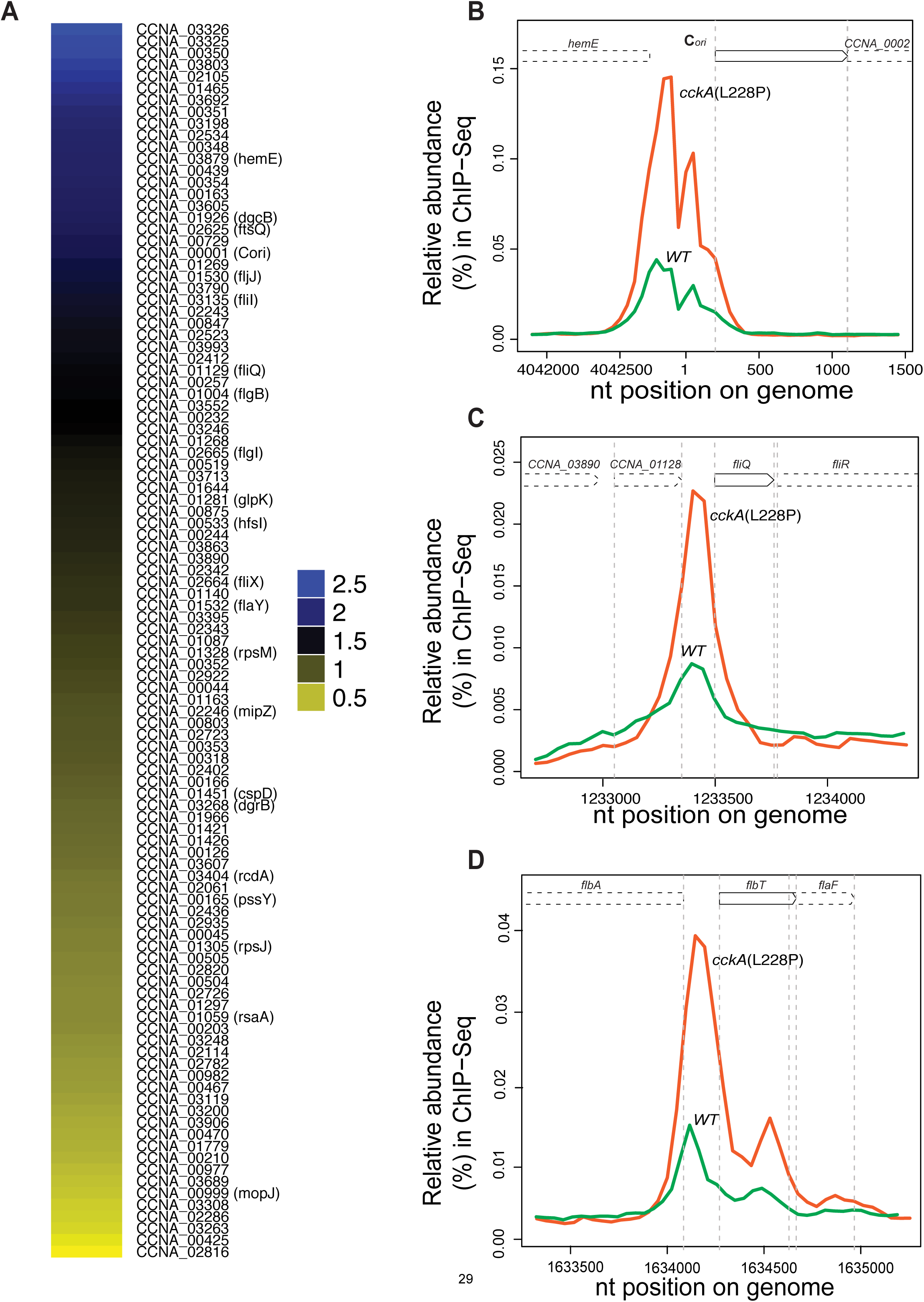
The CckA(L228P) mutation leads to differential CtrA binding. (A) Genome-wide comparative ChIP-Seq using polyclonal antibodies to CtrA, denoting the occupancy of CtrA on the chromatin of *cckA* vs ∆*nstA cckA*(L228P) mutant cells. The color key indicate the degree by which the occupancy of CtrA is varied in selected targets, as a result of the CckA(L228P) substitution. The color key at the bottom is expressed as log_2_ ratio (see the Supplementary data set for the complete list of target genes). Traces of the occupancy of CtrA at (B) the chromosomal origin of replication, C_*ori*_ (C) the promoter of *fliQ* (D) the promoter of *flbT.* The Figures 3B-D were derived from the ChIP-Seq data and the traces of CtrA in *WT* are denoted in green, and in ∆*nstA cckA*(L228P) mutant is denoted in orange.

Interestingly, in addition to the S-phase specific promoters, the ChIP-seq data also revealed a significant increase in binding of CtrA to the chromosomal origin of replication, C_*ori*_ (Figure 3B). The, qChIP experiment also confirmed the increase in CtrA occupancy at the C_*ori*_, in the *cckA*(L228P) mutant. In comparison to the *WT or* ∆*nstA*, a three-fold increase in the binding of CtrA at C_*ori*_ was evident in the *cckA*(L228P) mutant background (Supplementary Figure S3D). Together, these data pointed towards the differential DNA binding of CtrA, as a result of the *cckA*(L228P) point mutation to S-phase specific target promoters, and C_*ori*_, possibly regulated through the PAS-B domain of CckA.

### CtrA mediated repression of replication initiation mitigate NstA induced lethality

Next, we wondered if the increase in the CtrA binding to the C_*orI*_ is what contributes to the rescue of the toxicity induced by the overproduction of stable NstADD. We hypothesized that the specific modulation of CtrA activity by the CckA(L228P) mutation leading to enhanced binding of CtrA to the C_*ori*_ may be delaying replication initiation. Further, this delay in replication initiation may be slowing down the chromosome cycle to compensate for a reduced activity of a downstream event in the chromosome cycle. For example, the slow down in segregation process by the reduction in the decatenation activity of Topo IV by NstADD (18). To test this hypothesis, we decided to monitor the appearance or movement of the newly replicated chromosomal origin in the *cckA*(L228P) mutant. In *Caulobacter,* it has been well demonstrated that the newly formed origin of replication is immediately tethered to the opposite pole upon initiation of replication (29). The chromosome partitioning protein, ParB, specifically binds to the regions near C_*ori*_ and moves along with the C_*ori*_ upon initiation of replication (30,31). Therefore, the movement of fluorescently tagged ParB, GFP-ParB, can be used as a proxy to monitor the movement of the newly formed C_*ori*_ to the opposite cell pole (31,32). Localization experiments using GFP-ParB showed that the ∆*nstA cckA*(L228P) mutant had stalked cells with either single GFP-ParB foci (22.4%; Figure 4A red arrow heads, Figure 4C), or with two GFP-ParB foci, partially segregated, with the second foci still migrating to the opposite pole (33.8%; Figure 4A white arrow heads, Figure 4C). This was unlike in ∆*nstA* cells wherein the newly replicated C_*ori*_ along with GFP-ParB was immediately tethered to the opposite pole upon initiation of replication (86.9%; Figures 4B and C). From this observation we inferred that in the *cckA*(L228P) mutant, the initiation of replication and the elongation processes of the chromosome was slowed. This slow down may well be due to the increase in CtrA binding to the C_*ori*_. We also observed multiple GFP-ParB foci in *Caulobacter* cells, overproducing NstADD (Figure 4D), unlike the control samples with pMT335 vector alone, wherein, bipolar GFP-parB foci were predominant (Figure 4D). Thus we surmise that in the cells overproducing NstADD, multiple rounds of DNA replication are initiated and the chromosome decatenation is hampered (18). To counter this effect, *cckA*(L228P) point mutation enhances the CtrA binding at the C_*ori*_, which can significantly slow down the replication cycle. Our hypothesis was further corroborated by the observation that the increase in the binding of CtrA to the C_*ori*_ is still retained after the overexpression of NstADD. In comparison to the *WT* or ∆*nstA*, overexpressing NstADD, the *cckA*(L228P) mutant cells overproducing NstADD, had a significant increase in the occupancy of CtrA at the C_*ori*_ (Figure 5B).

**Figure 4.**
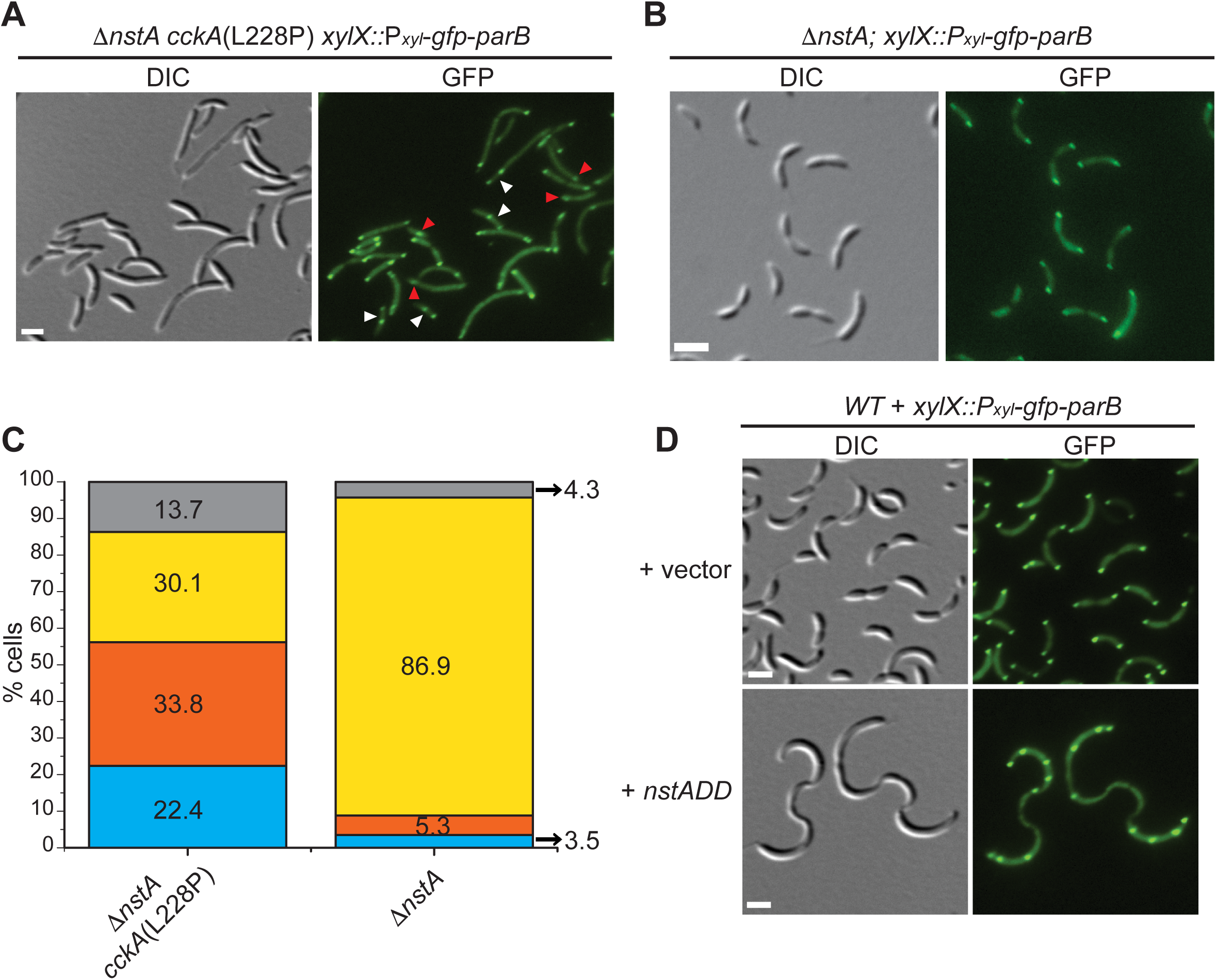
Suppression of NstADD toxicity by the *cckA*(L228P) mutant. (A) DIC and fluorescent micrographs of the extragenic suppressor mutant, ∆*nstA cckA*(L228P), harboring *gfp-parB* at the chromosomal *xylX* locus (xylX::P_*xyl*_-*gfp*-*parB*). Red arrow-heads: cells with one GFP-ParB foci; white arrow-heads: cells with two partially segregated GFP-ParB foci. (B) DIC and fluorescent micrographs of ∆*nstA* cells expressing *gfp-parB* from xylX::P_*xyl*_-*gfp*-*parB*. The cells were treated in the same manner, as described in A. (C) Data representing the stalked cells with one (blue), two partially segregated (orange), normal bipolar (yellow), and multiple (grey) GFP-ParB foci in ∆*nstA cckA*(L228P) (data from 1032 stalked cells) or ∆*nstA* (data from 944 stalked cells). (D) DIC and fluorescent micrographs of *WT* cells harboring *xylX*::P_*xyl*_-*gfp*-*parB*, and overexpressing NstADD from P_*van*_ on pMT335 or carrying the vector alone. The cells in (A), (B) and (D) were treated with 0.3% xylose to induce the production of GFP-ParB. Cells in (D) were additionally treated with 0.5 mM vanillate for 3h to induce NstADD production. Scale bar in A,B and D: 2µm.

**Figure 5.**
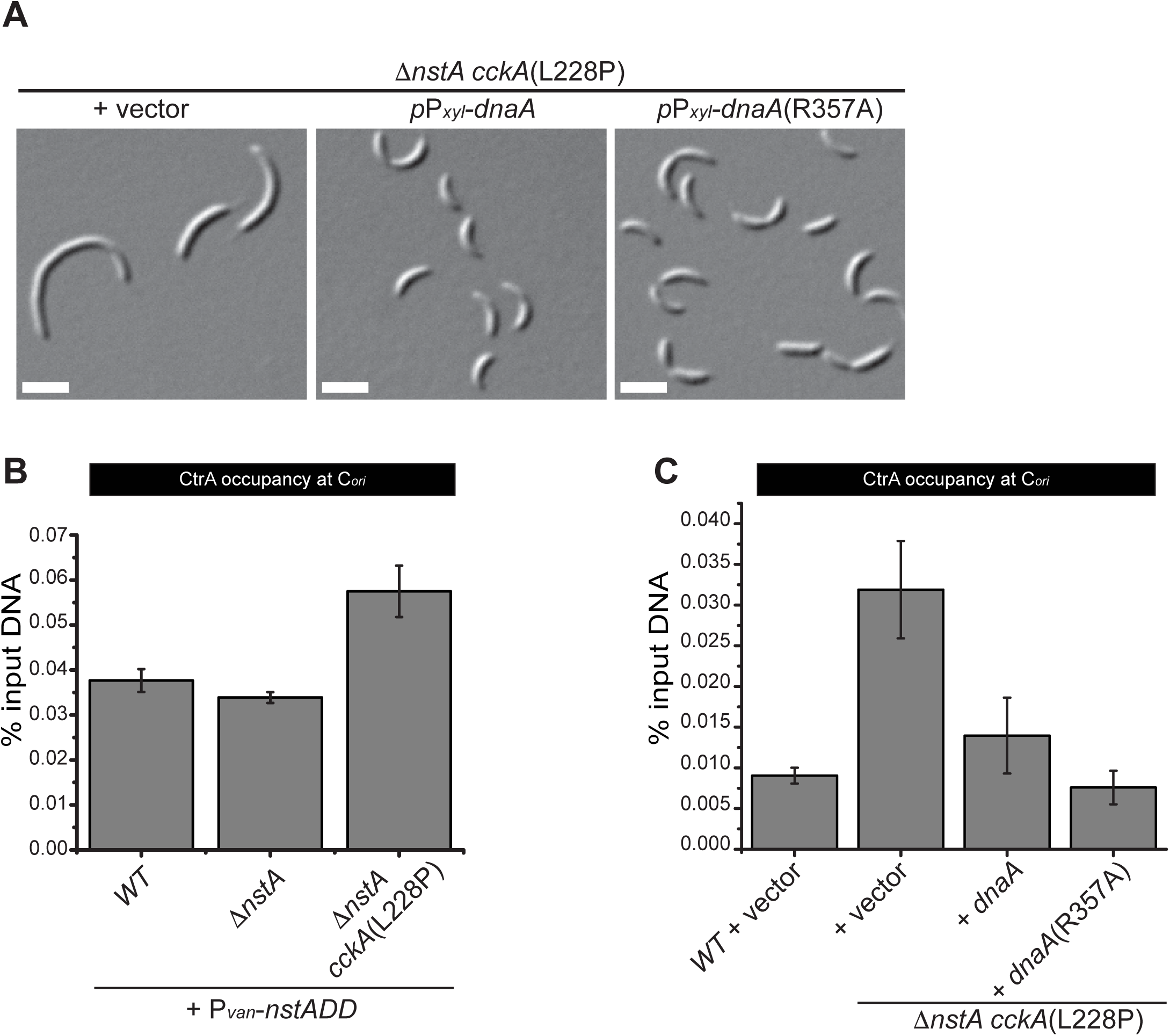
DnaA overexpression can alleviate the filamentation phenotype induced by the CckA(L228P). (A) DIC images showing ∆*nstA cckA*(L228P) harboring the vector or expressing *dnaA* or *dnaA*(R357A) from xylose inducible promoter on the medium copy vector, pJS 14. The cells were grown to exponential phase in PYE supplemented with 0.2% glucose, prior to the addition of 0.3% xylose. The xylose induction was done for 6h. (B) qChIP data depicting the CtrA occupancy at C_*ori*_ in *WT*, ∆*nstA*, and ∆*nstA cckA*(L228P) genetic backgrounds, after the overexpression of NstADD for three hours with 0.5mM vanillate inducer. (C) The qChIP data showing the CtrA occupancy at C_*ori*_ in ∆*nstA cckA*(L228P) cells, after the overexpression of DnaA or DnaA(R357A) from pJS14. The cells were treated in the same manner, as described in (A). The value ± SE, represented in (B) and (C) are the average of at least three independent experiments.

The CtrA binding boxes at the origin overlaps with the DnaA binding sites (6,33). Thus when CtrA∼P is abundant in the system, the DnaA binding to the origin is inhibited leading to inhibition of chromosome replication initiation (34). Therefore, we speculated that, if it is the increase in CtrA binding that is leading to slow down in replication, then such an inhibition should be relieved by titrating out CtrA at the C_*ori*_ by the overexpression of DnaA. Indeed, the overproduction of DnaA or its constitutively active ATP bound form, DnaA(R357A), from the xylose inducible promoter (P_*xyl*_) on a medium copy plasmid (35) caused a considerable decrease in cell filamentation of SN208 cells (Figure 5A). In addition, the qChIP experiments, also confirmed that the CtrA occupancy at C_*ori*_ was greatly reduced by about 70%, post the DnaA/DnaA(R357A) overexpression in the SN208 mutant (Figure 5C). Altogether, these results indicated at the increased binding of CtrA to C_*ori*_ facilitated by the *cckA*(L228P) mutation can alleviate toxicity attributed to NstADD overproduction.

## Discussion

The highly conserved CckA-CtrA signal transduction pathway in α-proteobacteria has several implications in development and pathogenesis. For example, during the early stages of symbiosis, in the nitrogen fixing bacteria, *Sinorhizobium meliloti* (*S. meliloti*), the role of CckA and its regulation has been shown to be essential (36). Likewise, the viability of the intracellular pathogen, *Brucella abortus* (*B. abortus*), in the human macrophages is dependent on the CckA-ChpT-CtrA pathway (37). The regulatory networks involving CtrA can be related to the specific lifestyle of the bacterium. For instance, while in *Caulobacter* CtrA is involved in cell-fate control, and cell cycle, by fine-tuning the stalked and swarmer cell programs, the control of cell envelope composition by CtrA is prevalent in *B. abortus* and *Rhizobium leguminosarum* (38,39), reiterating the plasticity of CckA-CtrA pathway.

In this study, we shown that the L228P mutation in the PAS-B domain of CckA not only increases the CtrA∼P levels but also rewires the preferential binding of CtrA to its target promoters (Figure 2A, 3 and Supplementary Dataset 1). The PAS-B domain in CckA has been shown to be necessary for regulation of its auto kinase activity, and for the switching of CckA between the kinase and the phosphatase modes (40). The CckA phosphatase activity during the swarmer to stalked cell transition is triggered by the binding of the effector molecule, c-di-GMP, to the PAS-B domain (16,17,40). Therefore, it is conceivable that the *cckA*(L228P) mutation possibly perturbs the binding of c-di-GMP to CckA thereby locking CckA in a kinase active form leading to increased CtrA∼P levels in the *cckA*(L228P) mutant (Figure 6).

**Figure 6.**
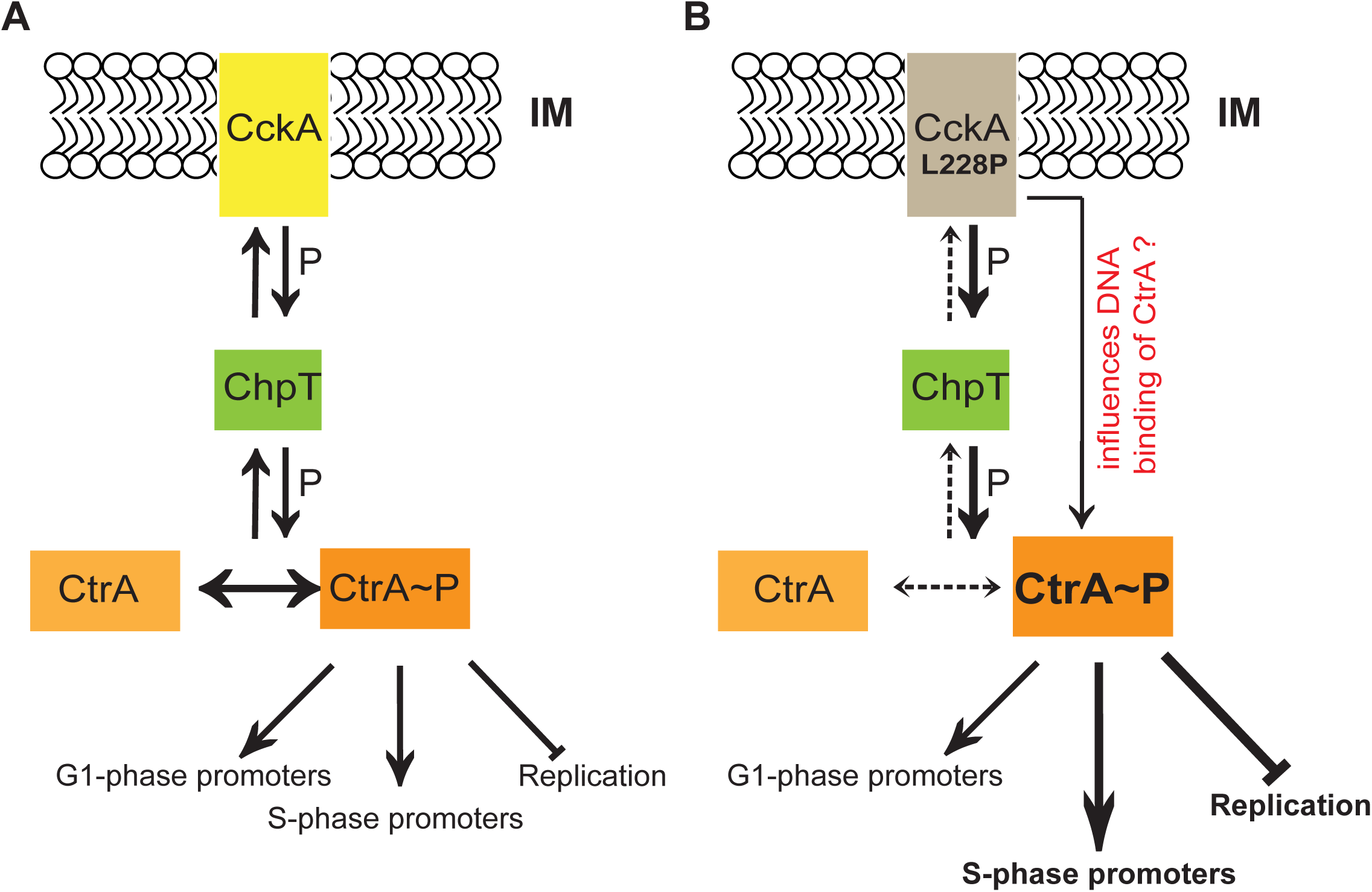
Model for CckA(L228P) function. (A) The membrane bound bi-functional kinase/phosphatase CckA (yellow), in its kinase form, phosphorylates CtrA (orange), via the intermediate phosphotransferase, ChpT (green). The active phosphorylated form of CtrA (CtrA∼P) binds to the DNA and triggers the transcription during G1- and late S-phase of the cell cycle, and inhibits chromosome replication in the G1 cells by binding to C_*ori*_. When the phosphatase activity of CckA is predominant (See Fig. 1A), the phosphate flow reverses dephosphorylating and inactivating CtrA. (B) The CckA(L228P) mutation possibly leads to a predominant CckA kinase acitivity thereby increasing CtrA∼P levels. In addition, the CckA(L228P) mutation, through an yet to be understood mechanism, specifically increases the CtrA∼P binding at C_*ori*_ and S-phase specific promoters.

Surprisingly, the above-mentioned increase in the CtrA∼P levels does not translate into a uniform increase in the binding of CtrA on all its target sites on the chromosome. The increased binding happens only at the C_*ori*_ and a sub-set of S-phase specific CtrA promoters. Previous studies have shed light on the additional components, SciP and MucR, which modulate the activity of the CtrA-dependent promoters during cell cycle (41–43). While MucR specifically represses G1-phase promoters of CtrA, SciP has been shown to negatively regulate the CtrA-dependent promoters whose activity are known to peak at the S-phase of the cell cycle (42). Interestingly, our comparative ChIP-Seq analysis, revealed that the CckA(L228P) substitution contributes to specific enhancement of CtrA binding at S-phase promoters, which are also bound by SciP. The mechanistic mode of repressing the CtrA transcription by SciP involves a direct interaction between SciP and CtrA (43). Interestingly, SciP does not perturb the DNA binding activity of CtrA and it blocks the RNA polymerase recruitment to the CtrA activated promoters (43). Moreover, SciP itself is under the direct transcriptional control of CtrA (41). Nevertheless, our ChIP-Seq analysis shows that in the *cckA*(L228P) mutant the binding of CtrA to the *sciP* promoter is not significantly altered (Supplementary Dataset 1) indicating that *sciP* transcription might not be altered in CckA(L228P). Therefore, it is tempting to speculate that the CckA(L228P) substitution possibly facilitates CtrA to overcome the inhibition imparted by SciP either by directly acting on SciP or by making CtrA more potent to compete for the RNA polymerase. It may also be possible that the CckA kinase could be regulating the interaction between SciP and CtrA, in a direct or indirect manner. However, this hypothesis, remains to be investigated further and our results pave way for exploring this intriguing aspect of the CckA-CtrA pathway.

## Materials and methods

### Growth Conditions and Media

*Caulobacter* strains were grown on rich PYE media (0.2% peptone, 0.1% yeast extract, 1 mM MgSO_4_, 0.5 mM CaCl_2_) or minimal M2G media (M2 -1X salt solution [Na_2_HPO_4_ 0.87gm/L, KH_2_PO_4_ 0.53gm/L, NH_4_Cl 0.25gm/L] supplemented with 0.5 mM MgSO_4_, 0.2 % Glucose, 10 µM FeSO_4_.EDTA, 0.5 mM CaCl_2_) (44), and incubated at 29°C, unless specifically mentioned. The *Caulobacter* strains were subjected to electroporation, øCr30 mediated transductions, and intergeneric conjugations (using *E.coli S17-1*) as previously described (24,45,46). *E. coli* strains, EC100D (Epicentre, WI, USA), and S17-1 were grown on LB media and incubated at 37°C, unless specifically mentioned.

### In vivo phosphorylation

In vivo phosphorylation experiments were performed as described previously (47). A single colony of cells picked from a PYE agarose plate was washed with M5G medium lacking phosphate and was grown overnight in M5G with 0.05 mM phosphate to an optical density of 0.3 at 660 nm. One milliliter of culture was labeled for 4 min at 28°C using 30 µCi of γ-[^32^P]ATP. Upon lysis, proteins were immunoprecipitated with 3 ml of anti-CtrA antiserum and Protein A agarose (Roche, CH) and the precipitates were resolved by SDS–polyacrylamide gel electrophoresis and radiolabelled CtrA was quantified and were normalized to the relative cellular content as determined by immunoblotting of lysates.

### Extragenic suppressor Screen

*∆nstA* or *WT* cells were UV irradiated with 700 and 900×100 µJ/cm^2^ energy using CL 1000 UV Cross linker (UVP, Cambridge, UK). The irradiated cells were diluted into PYE media followed by 6 hr incubation. The cells were then electroporated with plasmid pMT335-P_*van*_-*nstA*DD and plated on PYE supplemented with gentamycin and vanillate inducer. Individual colonies were then grown in liquid PYE containing gentamycin, and vanillate inducer for overnight. Two criteria were used for confirming the extragenic mutation: (i) plasmids from the selected mutants were again transformed into *WT Caulobacter* and checked for *nstA* toxicity, to avoid the possibility that the suppression is due to mutation on the plasmid, and (ii) plasmid cured mutants were retransformed with fresh pMT335-P_*van*_-*nstA*DD, to confirm that the toxicity suppression is indeed due to a mutation in the chromosome. Mutations were mapped by next generation sequencing on an Illumina platform at Fasteris, Switzerland.

### Chromatin immunoprecipitation (ChIP)

ChIP experiments were carried out as described earlier (24). Mid-log phase cells were cross-linked in 10 mM sodium phosphate (pH 7.6) and 1% formaldehyde at room temperature for 10 minutes and on ice for 30 min thereafter, washed thrice in phosphate buffered saline (pH 7.4) and lysed in 5000 Units of Ready-Lyse lysozyme solution (Epicentre Technologies, WI, USA). Lysates were sonicated on ice using 7 bursts of 30 sec to shear DNA fragments to an average length of 0.3-0.5 kbp. The cell debris were cleared by centrifugation at 14,000 rpm for 2 min at 4°C. Lysates were normalized by protein content, diluted to 1 mL using ChIP buffer (0.01% SDS, 1.1% Triton X-100, 1.2 mM EDTA, 16.7 mM Tris-HCl [pH 8.1], 167 mM NaCl plus protease inhibitors [Complete^TM^ EDTA-free, Roche, Switzerland]), and pre-cleared with 80 µl of protein-A agarose (Roche, Switzerland) saturated with 100 µg BSA and 300 µg Salmon sperm DNA. Ten % of the supernatant was removed and used as total chromatin input DNA. To the remaining supernatant, anti-CtrA (20) antibody was added (1:500 dilution), and incubated overnight at 4°C. Immuno complexes were trapped with 80 µL of protein-A agarose beads pre-saturated with BSA-salmon sperm DNA. The beads were then washed once each with low salt buffer (0.1% SDS, 1% Triton X- 100, 2 mM EDTA, 20 mM Tris-HCl [pH 8.1], 150 mM NaCl), high salt buffer (0.1% SDS, 1% Triton X-100, 2 mM EDTA, 20 mM Tris-HCl [pH 8.1], 500 mM NaCl) and LiCl buffer (25 mM LiCl, 1% NP-40, 1% sodium deoxycholate, 1 mM EDTA, 10 mM Tris-HCl [pH 8.1]), and twice with TE buffer (10 mM Tris-HCl[pH 8.1], 1 mM EDTA). The protein•DNA complexes were eluted in 500 µL freshly prepared elution reagent (1% SDS, 1 mM NaHCO_3_). This was supplemented with NaCl to a final concentration of 300 mM and incubated overnight at 65°C to reverse the crosslinks. The samples were treated with 2 µg of Proteinase K (Roche, Switzerland) for 2 hr at 45°C in after addition of 40 mM EDTA and 40 mMTris-HCl (pH 6.5). DNA was extracted using phenol:chloroform:isoamyl alcohol (25:24:1), ethanol-precipitated using 20 µg of glycogen as carrier, and resuspended in 50 µL of sterile deionized water. The comparative ChIP-followed by deep Sequencing (ChIP-Seq), was done using the next generation sequencing on an Illumina platform at Fasteris, Switzerland.

### Quantitative PCR (qPCR) analyses

qPCR was performed on a CFX96 Real Time PCR System (Bio-Rad, CA, USA) using 10% of each ChIP sample, 12.5 µL of SYBR^®^ green PCR master mix (Bio-Rad, CA, USA), 200 nM of primers and 6.5 µl of water per reaction. Standard curve generated from the cycle threshold (Ct) value of the serially diluted chromatin input was used to calculate the % input value of each sample. Average values are from triplicate measurements done per culture. The final data was generated from three independent cultures. The SEM shown in the figures was derived with Origin 7.5 software (OriginLab Corporation, Northhampton, MA, USA). C_*ori*__Fwd (5’-CGCGGAACGACCCACAAACT-3’) and C_*ori*__Rev (5’-CAGCCGACCGACCAGAGCCA-3’) primer pairs as described earlier (35) were used to amplify the region near *ori* precipitated by anti-CtrA antibody. To check CtrA binding on P_*pilA*_, the DNA region analysed by real time PCR was from nucleotide −287 to −91 relative to the start codon of *pilA* (24). A P_*kidO*_ fragment comprising nt 3,857,810–3,858,141 of the NA1000 genome sequence was quantified (23) to monitor CtrA binding on P_*kidO*_. To quantify CtrA occupancy at the promoter of *flbT*, the DNA region, −280 to +30 relative to the start codon of *flbT*, was used. The DNA region from −226 to +30 relative to the start codon of *tacA* was analysed for quantifying CtrA occupancy on P_*tacA*_ (24).

### ChIP-seq data analysis

The FASTQ files were checked for quality of sequencing using FastQC software, version 0.11.5. The first ten bases showed distortion, due to which it was decided to trim the first ten bases from all short reads. The reads were trimmed at the 5' end for 10 bases using fastx_trimmer tool from Fastx-toolkit version 0.0.14. The preprocessed reads were mapped to the *Caulobacter Cresentus* NA1000 reference genome (CP001340.1) using aligner Bowtie version 1.0.0 using the following parameter: -m 1, - S, -v 2. Around 36.9 million reads mapped uniquely to the reference genome for the wild type *cckA* and 21 million reads for the mutant *cckA*(L228P) strains.

Further, the aligned reads were imported onto Seqmonk (version 1.38.1) to build the sequence read profiles. The genome was subdivided into 50bp probes and a value representing the number of reads mapping to the genome within a probe was calculated using the Read Count Quantitation option. The probe list with the quantified value for each probe was exported. Custom Perl scripts were used to compute the relative abundance of each probe as a percent with respect to the total uniquely mapped reads for each dataset. A cutoff was determined as average reads plus twice the standard deviation of the sample to differentiate between candidate peaks and background noise. The candidate peaks were annotated using custom Perl scripts. A probe was annotated with a gene if the centre of the probe was within a distance of −500 and +100 bases from the transcription start site of the gene, taking into account the orientation of the gene as well. If a probe is found to satisfy the condition for two genes (each on either strand), then both the genes are reported. A list of nearby RNA genes are also reported separately. Probes without an annotation are labeled as ‘NO ANNO’.

### Microscopy

Differntial interference contrast (DIC) and fluorescence microscopy were performed on a Nikon Eclipse 90i microscope equipped with 100X oil TIRF (1.49 numerical aperture) objective and a coolSNAP HQ-2 (Photometrics, USA) CCD camera. Cells were placed on a 1% agarose solidified pads for imaging. Images were processed and analyzed with the Metamorph software (Molecular Devices, USA).

### β-Galactosidase Assay

The cultures harbouring were incubated at 29°C till it reached 0.1-0.4 OD@660 nm (A_660_). 50 µl of the cells were treated with a 10 µl of chloroform followed by the addition of 750 µ l of Z-buffer (60 mM Na_2_HPO_4_, 40 mM NaH_2_PO_4_, 10 mM KCl, 1mM MgSO_4._7H_2_O, pH 7.0) followed by 200 µl of Ortho Nitro Phenyl-β-D-Galactoside (from stock concentration of 4 mg/ml dissolved in 100 mM potassium phosphate buffer [pH 7.0]). The reaction mixture was incubated at 30°C till yellow color was developed. Finally 500µl of 1 M Na_2_CO_3_ solution was added to stop the reaction and absorbance at 420nm (A_420_) of the supernatant was noted using Z-buffer as the blank. The miller units (U) were calculated using the equation U= (A_420_ X 1000) / (A_660_ X t X v), where ‘t’ is the incubation time (min), ‘v’ is the volume of culture taken (ml). Experimental values were average of three independent experiments. The SEM shown in the figures was derived with Origin 7.5 software (OriginLab Corporation, Northhampton, MA, USA).

## Acknowlegdements

We thank Justine Collier and Patrick Viollier for materials, Rashmi Sukumaran for her assistance with the ChIP-seq analysis, and Srinivasa Murty Srinivasula for discussions and critical comments on the manuscript. S.N. is supported by a graduate fellowship from IISER Thiruvananthapuram. L.K is supported by a Junior Research Fellowship from the Council of Scientific & Industrial Research (CSIR) India. This work was supported by funds from the Wellcome Trust-DBT India Alliance (500140/Z/09/Z) through an Intermediate fellowship to S.K.R., and funds from the SwarnaJayanti Fellowship (DST/SJF/LSA-01) from Department of Science and Technology, India, to S.K.R.

## SUPPLEMENTARY INFORMATION

**Supplementary Figure S1.**
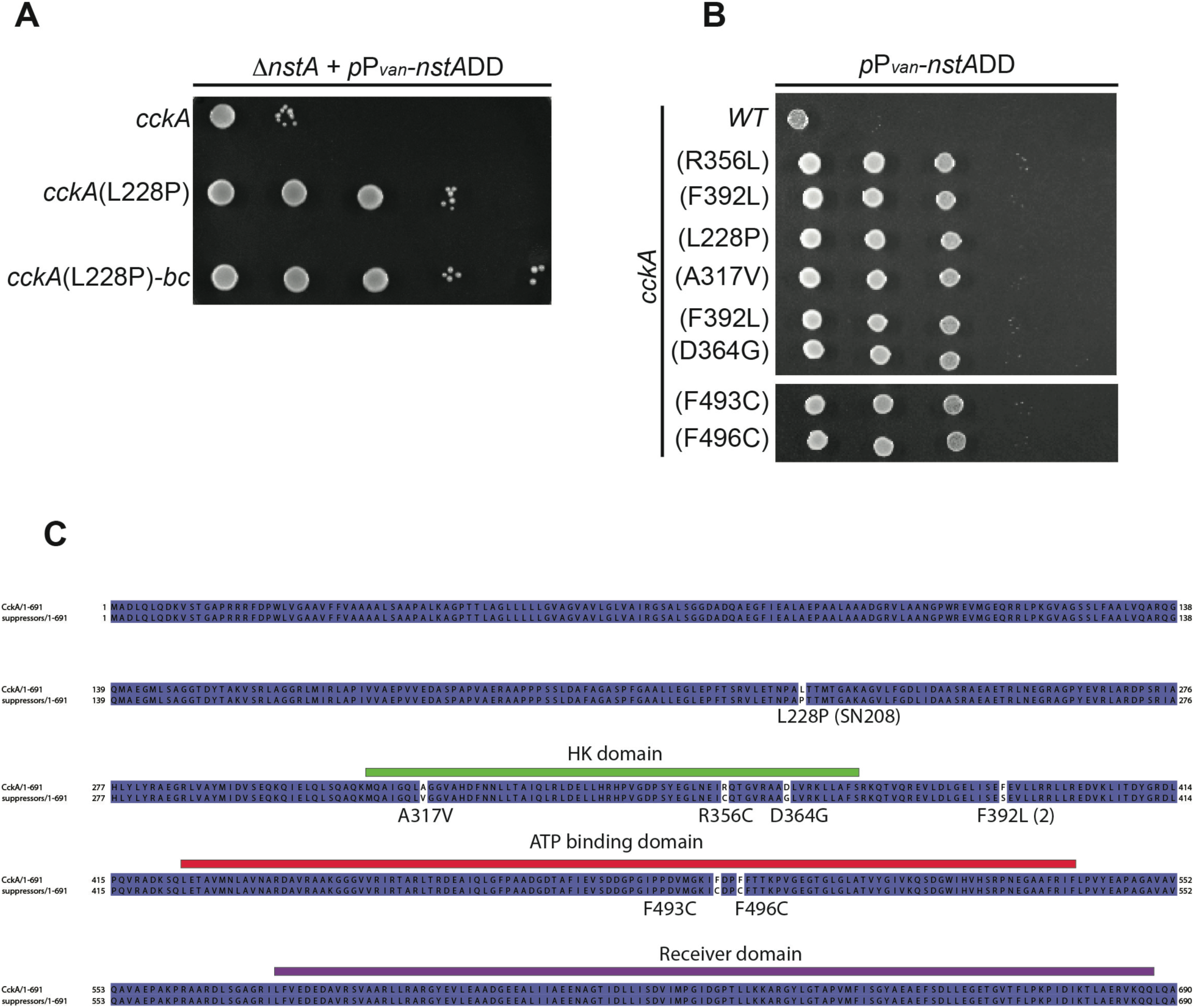
(A) Growth of ∆*nstA* cells overproducing NstADD, and harboring either *wild-type cckA* or *cckA*(L228P) or *cckA*(L228P)-back-cross (bc). (B) Growth of *Caulobacter* harboring the *wild-type cckA*, or NstADD toxicity suppressor mutants of *cckA*, upon *nstA*DD overexpression. In (A) and (B), cells were diluted fivefold and spotted on media containing 0.5 mM vanillate. The mutant *cckA*(F496C) is in the ∆*nstA* genetic background, the rest were in *WT* background. (C) Schematic denoting the CckA suppressor mutations. The CckA(L228P) mutation resides in the PAS-B domain, whereas the other point mutations are harbored either in the histidine kinase domain or the ATP binding domain of CckA.

**Supplementary Figure S2.**
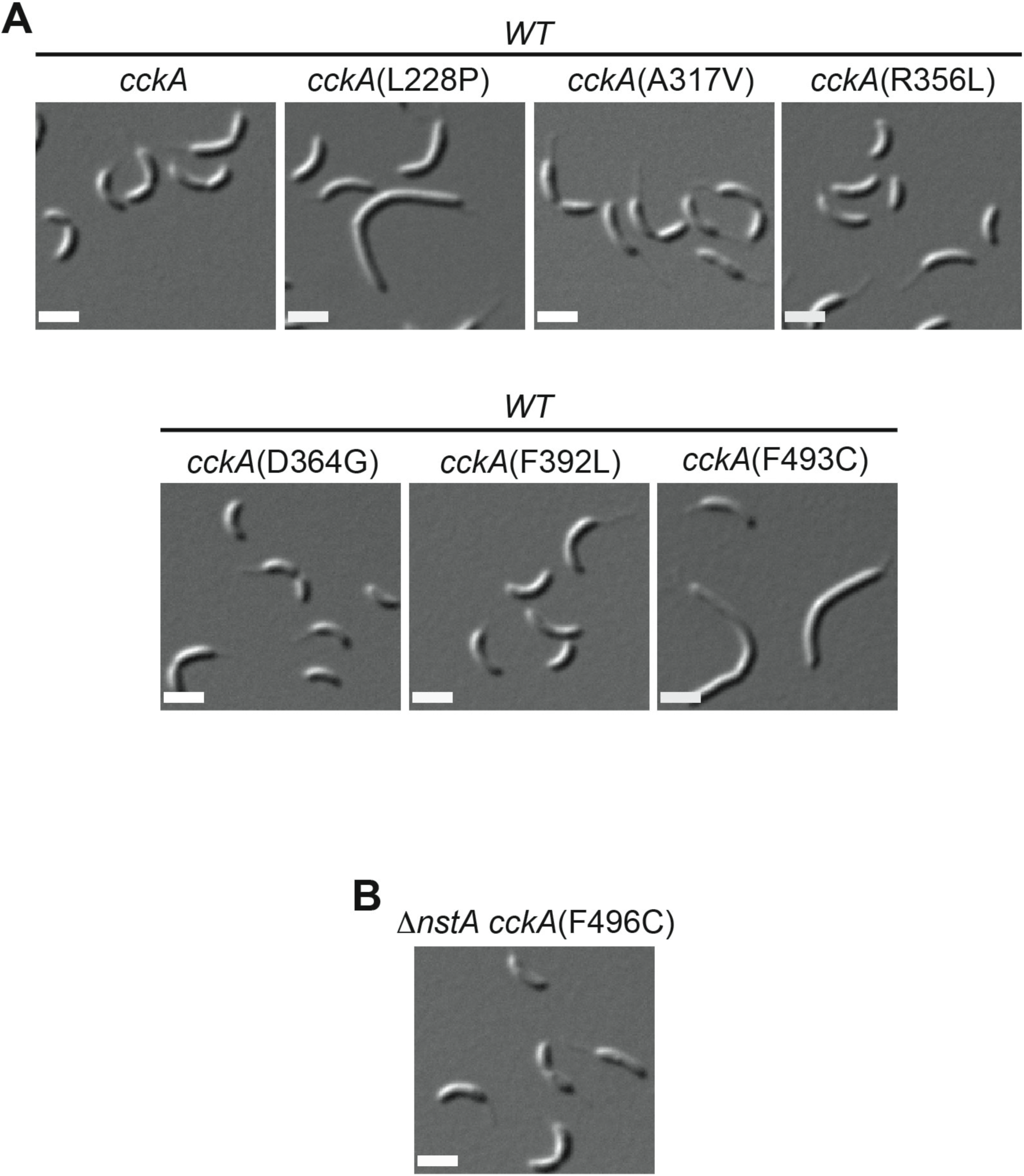
(A) DIC images of the various point mutants of *cckA* in *WT* genetic background. (B) DIC image of ∆*nstA cckA*(F496C) mutant.

**Supplementary Figure S3.**
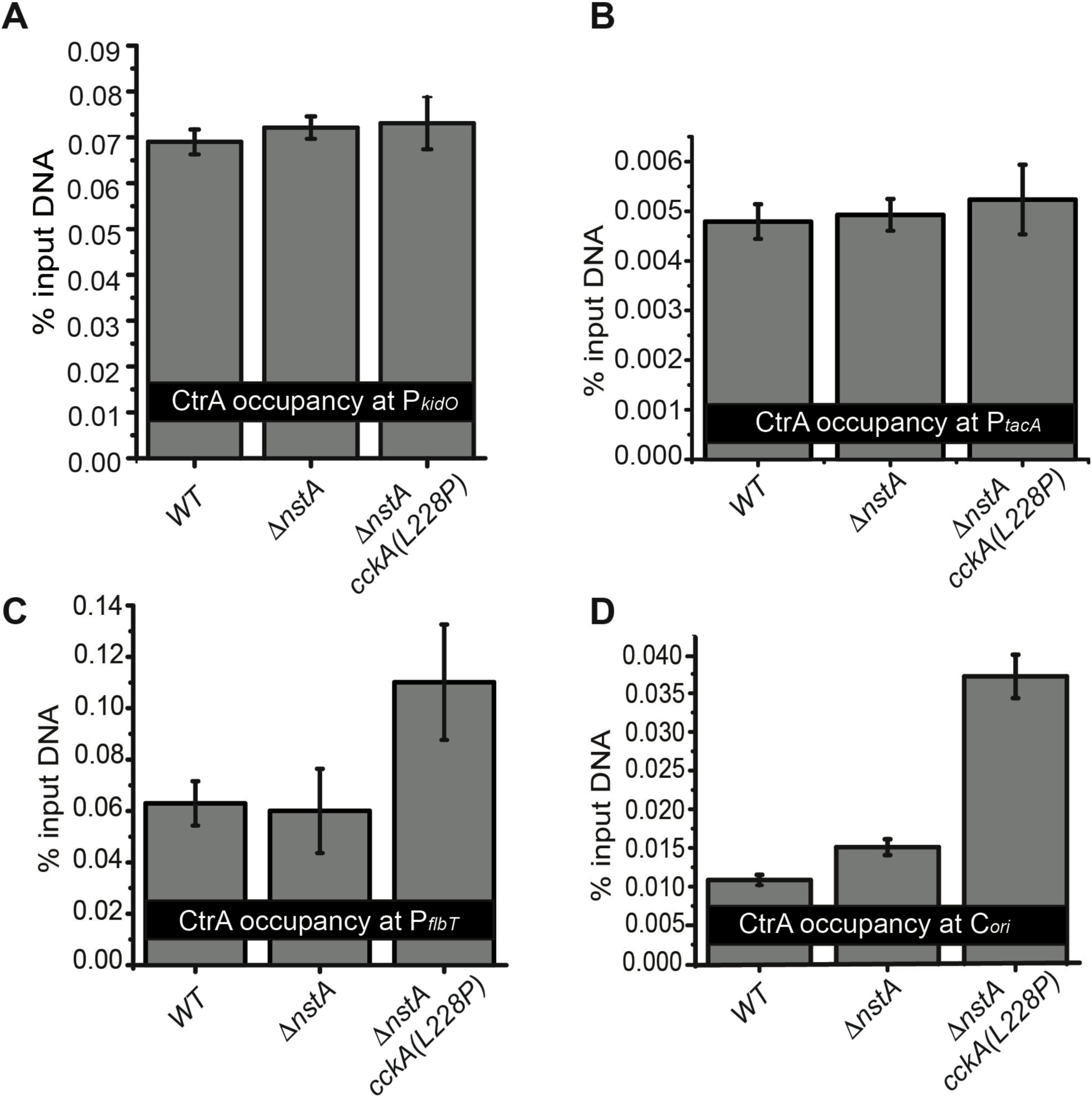
The qChIP data indicating the CtrA occupancy at (A) the promoter of *kidO* (P_*kidO*_), (B) the promoter of *tacA* (P_*tacA*_), (C) the promoter of *flbT* (P_*flbT*_) and (D) at the chromosomal origin of replication, C_*ori*_, in *WT,* ∆*nstA*, and ∆*nstA cckA*(L228P) cells. The data represented are the average of 3 independent experiments ± SE.

**Supplementary Figure S4.**
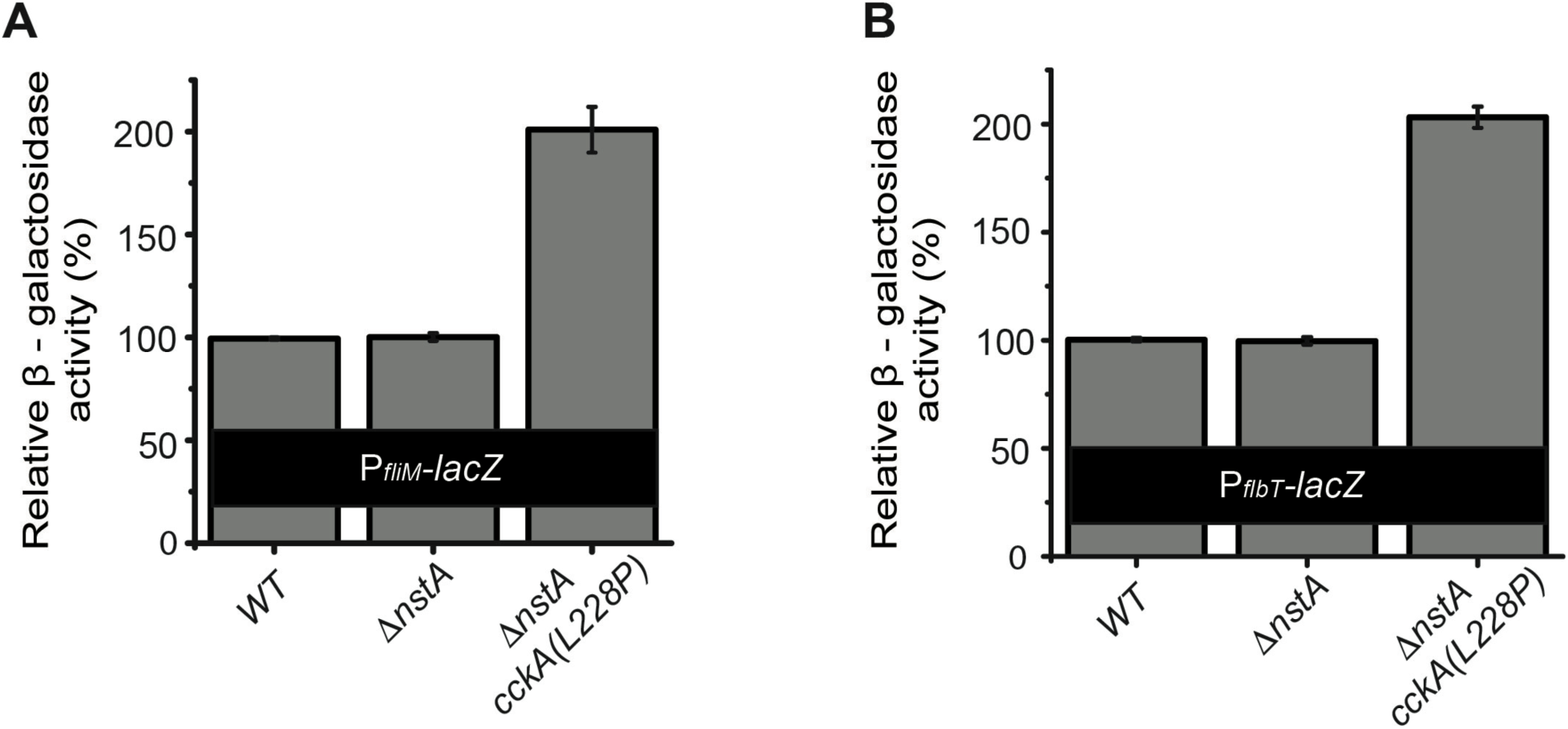
Relative β-galactosidase activities of (A) P_fliM_-*lacZ* reporter, and (B) P_*flbT*_-*lacZ* reporter, in *WT*, ∆*nstA*, and ∆*nstA cckA*(L228P) cells. The data represented in (A) and (B) are the average of 3 independent experiments ± SE.

## Supplementary Methods

### Strain construction

The various *cckA* point mutant strains namely **SN208** [*∆nstA cckA* (L228P)], **LK98** [*WT cckA*(R356C)], **LK100** [*∆nstA cckA*(F496C)], **LK109** [*WT cckA*(L228P)], **LK122** [*WT cckA*(A317V)], **LK124** [*WT cckA*(F493C)], **LK128** [*WT cckA*(F392L)] and **LK135** [*WT cckA*(D364G)] were generated by Ultraviolet (UV) radiation based mutagenesis using *WT* or SKR1797 (*∆nstA*) (1).

The strains namely **SN227** [*∆nstA cckA* (L228P) + pMT 335-*nstA*DD], **SN1140** [*WT cckA*(L228P) + pMT335-P_*van*_-*nstA*DD], **SN1141** [*WT cckA*(A317V) + pMT335-P_*van*_-*nstA*DD], **SN1142** [*WT cckA*(F493C) + pMT335-P_*van*_-*nstA*DD], **SN1144** [*WT cckA*(F392L) + pMT335-P_*van*_-*nstA*DD], **SN1145** [*WT cckA*(D364G) + pMT335-P_*van*_-*nstA*DD], **SN1151** [*WT cckA*(R356C) + pMT335-P_*van*_-*nstA*DD] and **SN1152** [*∆nstA cckA*(F496C) + pMT335-P_*van*_-*nstA*DD] were made by electroporating pSKR126 (*pP_van_-nstA*DD) (1) into SN208 [*∆nstA cckA* (L228P)], LK109 [*WT cckA*(L228P)], LK122 [*WT cckA*(A317V)], LK124 [*WT cckA*(F493C)], LK128 [*WT cckA*(F392L)], LK135 [*WT cckA*(D364G)], LK98 [*WT cckA*(R356C)] and LK100 [*∆nstA cckA*(F496C)] respectively,

The strains **SN377** (*WT; xylX*::P_*xyl*_-*gfp-parB*),**SN379** [*∆nstA cckA*(L228P); *xylX::P_xyl_-gfp-parB*] and **SN559** (*∆nstA; xylX*::P_*xyl*_-*gfp-parB*) were made by electroporating pSN190 (pXGFP4C1-P_*xyl*_-*gfp-parB*) into *WT,* SN208 and SKR1797 respectively.

The strains **SN461** [*∆nstA cckA*(L228P) + pJSX-*dnaA*], **SN465** [*∆nstA cckA*(L228P) + pJSX-*dnaA* (R357A)] and **SN467** [*∆nstA cckA*(L228P)+pJS14] were made by electroporating pJSX-*dnaA*, pJSX-*dnaA* (R357A) (2) and pJS14 into SN208, respectively.

The strains **SN505** (*WT*; *xylX::* P_*xyl*_-*gfp-parB* + pMT335) and **SN1153** ( *WT*; *xylX::*P_*xyl*_-gfp-parB + pMT335-P_*van*_-*nstA*DD) were made by electroporating pMT335 (3) and pSKR126 into the strain SN377, respectively.

The strains **SN740** (*WT* + pLac290-P_*pilA*_-*lacZ*), **SN742** (*∆nstA* + pLac290-P_*pilA*_-*lacZ*) and **SN744** [*∆nstA cckA*(L228P) + pLac290-P_*pilA*_-*lacZ*) were made by electroporating pJS70 (pLac290-P_*pilA*_-*lacZ*) (4) into *WT,* SKR1797 and SN208, respectively.

The strains **SN741** (*WT* + pLac290-P_*tacA*_-*lacZ*), **SN743** (*∆nstA* + pLac290-P_*tacA*_-*lacZ*) and **SN745** [*∆nstA cckA*(L228P) + pLac290-P_*tacA*_-*lacZ*) were made by electroporating pMV05 (pLac290-P_*tacA*_-*lacZ*) (5) into *WT,* SKR1797 and SN208, respectively.

The strains **SN1361** (*WT* + pLac290-P_*fliM*_-*lacZ*), **SN1365** (*∆nstA* + pLac290-P_*fliM*_-*lacZ*) and **SN1369** [*∆nstA cckA*(L228P) + pLac290-P_*fliM*_-*lacZ*) were made by electroporating pLac290-P_*fliM*_-*lacZ* into *WT,* SKR1797 and SN208, respectively.

The strains **SN1406** (*WT* + pLac290-P_*flbT*_-*lacZ*), **SN1407** (*∆nstA* + pLac290-P_*flbT*-_*lacZ*) and **SN1408** [*∆nstA cckA*(L228P) + pLac290-P_*flbT*_-*lacZ*) were made by electroporating pLac290-P_*flbT*_-*lacZ* into *WT,* SKR1797 and SN208, respectively.

The *cckA*(L228P) back cross strain **SN769** [*∆nstA*; *cckA*(L228P)] was made by backcrossing the *cckA*(L228P) point mutation in SN208 into SKR1797. pSN155 (pNPTS-*cckA* backcross construct) was used to transform SN208 and the transformants were plated on PYE supplemented with Kanamycin. Further øCr30 lysates of the transformants were made and was used for transducing into SKR1797 thereby generating SN769. The backcross strain, SN769, was electroporated with pSKR126, to obtain **SN771** [*∆nstA*; *cckA*(L228P) + pMT335-P_*van*_-*nstA*DD].

The strain SKR1800 (*WT* + pMT335-P_*van*_-*nstA*DD) is previously described (1).

### Plasmid construction

The plasmid **pSN155** (pNPTS-*cckA*-backcross) construct was made by PCR amplifying a region 750bp (approx.) upstream of *cckA*. The PCR fragment was digested with *Eco*RI/*Hind*III. The digested fragment was ligated into pNPTS138 (M.R.K Alley, unpublished) cut with *Eco*RI/*Hind*III.

Plasmid **pSN190** (pXGFP4-C1-P_xyl_-*gfp-parB*) was made by PCR amplifying *parB* and cleaving it with *BgI*II/*Eco*RI, wherein the predicted the start codon ATG was replaced with GTG that carried an overlapping *Bgl*II recognition site to allow proper placement of *parB* facilitating N-terminal GFP fusion. The alleles were ligated into pXGFP4-C1 vector (M.R.K Alley unpublished) cut with *Bgl*II/*Eco*RI.

To make **pSN205** (P_*fliM*_-*lacZ*, a kind gift from Patrick Viollier) nucleotides 2298862-2299971 of NA1000 genome (CP001340) was amplified and ligated as *Eco*RI/*Hind*III fragment into a medium-copy plasmid pJGZ290 (6) to drive the transcription of the promoterless *lacZ* gene.

Plasmid **pSN206** (P_*flbT*_-*lacZ*, a kind gift from Patrick Viollier) was made by amplifying nucleotides 1633750-1634327 of the NA1000 genome (CP001340) and ligating as *Eco*RI/*Hind*III fragment into the medium-copy plasmid pJGZ290 (6) to drive the transcription of the promoterless *lacZ* gene.

